# Impact of within-tree organ distances on floral induction and fruit growth in apple tree: implication of carbon and gibberellin organ contents

**DOI:** 10.1101/612317

**Authors:** Farés Belhassine, Sébastien Martinez, Sylvie Bluy, Damien Fumey, Jean-Jacques Kelner, Evelyne Costes, Benoît Pallas

## Abstract

In plants, organs are inter-dependent for growth and development. Here, we aimed to investigate the distance at which interaction between organs operates and the relative contribution of within-tree variation in carbohydrate and hormonal contents on floral induction and fruit growth, in a fruit tree case study. Manipulations of leaf and fruit numbers were performed in two years on ‘Golden delicious’ apple trees, at the shoot or branch scale or one side of Y-shape trees. For each treatment, floral induction proportion and mean fruit weight were recorded. Gibberellins content in shoot apical meristems, photosynthesis, and non-structural carbohydrate concentrations in organs were measured. Floral induction was promoted by leaf presence and fruit absence but was not associated with non-structural content in meristems. This suggests a combined action of promoting and inhibiting signals originating from leaves and fruit, and involving gibberellins. Nevertheless, these signals act at short distance only since leaf or fruit presence at long distances had no effect on floral induction. Conversely, fruit growth was affected by leaf presence even at long distances when sink demands were imbalanced within the tree, suggesting long distance transport of carbohydrates. We thus clarified the inter-dependence and distance effect among organs, therefore their degree of autonomy that appeared dependent on the process considered, floral induction or fruit growth.

## Introduction

In plants, the determination of organ nature, their development and growth are considered as interdependent. For instance, the position at which flowers develop is linked to the number of nodes developed from the seed (Sachs, 1999). Architectural analyses have highlighted highly structured organization in a large range of plants, with particular organ types being observed at particular positions and times during ontogeny (Barthélémy and Caraglio, 2007). This has been demonstrated for instance for the position of reproductive organs in *Quercus ilex* or *Pinus halepensis* (Barthélémy and Caraglio, 2007) or for flower buds along axes in different *Rosaceae* species (Costes *et al.*, 2014). The inter-dependence and differential development of organs within plants are assumed to depend on water, carbohydrates, hormones, mineral nutrients, etc. that are transported within the plants. Both the availability of these resources and the total number of competing organ define a developmental and growth context for each organ depending on its position during plant life span. Among these shared resources, carbohydrates have been particularly studied as they are considered as a main limiting factor for organ growth (Grossman and DeJong, 1995) and as a regulator of the transition between vegetative and reproductive phase in plant life (Rolland *et al.*, 2006). In the particular case of fruit trees, the number, position of fruits, as well as their growth at harvest are dependent on the capability of a given meristem to be floral, then of this flower to fruit set, and finally of a fruit to capture resources for its growth.

In fruit trees, the capability of a shoot apical meristems (SAM) to be floral induced is strongly affected by the presence of fruit during the growing season. A first hypothesis explaining floral induction (FI) inhibition in conditions of high crop load is associated with a competition for carbohydrates between meristems and fruit (Monselise and Goldschmidt, 1982). Besides this “carbon” hypothesis, Chan and Cain (1967) have demonstrated that FI is inhibited by seed development through hormones. This hypothesis was confirmed by experiments on seedless apple and pear cultivars suggesting that seeds may inhibit FI, probably by gibberellins (Dennis and Neilsen, 1999). Gibberellins (GA) are considered among the pathways involved in floral induction control in *Arabidopsis thaliana* (Jung *et al.*, 2017). However, their effect is currently considered as inverse in *Arabidopsis thaliana* and in perennial woody plants. Indeed, GA promotes the transition from vegetative to reproductive development of buds in Arabidopsis (Wilson *et al.*, 1992), while it is assumed to inhibit FI in fruit trees such as mango (Nakagawa *et al.*, 2012) and apple (Wilkie *et al.*, 2008). GA12 has been observed as the transported GA form moving within the plant through the vascular system in *Arabidopsis* (Regnault *et al.*, 2015). In the apple tree, GA4 has been assumed to move from fruit to SAM in apple tree (Ramírez *et al.*, 2004). The involvement of GA in FI control was further confirmed by differential expressions of genes involved in the GA biosynthesis pathway (GA20ox and GA2ox) in SAM of apple trees with heavy or low crop loads (Guitton *et al.*, 2016). This study also suggested a context of carbohydrate starvation, in SAM of trees in high cropping conditions. Therefore, the co-involvement of carbohydrate and hormones in FI control appears as an assumption to further investigate. It implies the involvement of several processes: photosynthesis by leaves, transport from leaves to sinks, including SAM, but also the presence of GA in SAM likely the active forms GA4 and GA1 (Ramírez et al., 2004). Moreover, leaves may have a dual role in FI control since, in addition to be source of carbohydrates, they are also producing FLOWERING LOCUS T (FT) protein, which is transported to the SAM to activate floral induction in many species, including fruit species (Hanke *et al.*, 2007). Nevertheless, it is currently still unclear at which distance the different “signals” originating from fruit and leaves act on FI in SAM.

Regarding carbohydrates, partitioning from sources to sinks is considered as a function of source supply, sink demand and distances between them (Lacointe 2000). Nevertheless, in fruit trees there is no clear consensus about the impact of distances between sources and sink on carbohydrate allocation and on their consequences on the existing organ growth variability within the trees. Depending on their strength, i.e. the ability of an organ to import assimilate, sinks can use carbohydrates from nearby or distant sources. Carbohydrates can move at short distances, i.e. from non-fruiting to fruiting shoots (Walcroft *et al.*, 2004; Pallas *et al.*, 2018) to sustain fruit growth or at longer distances, i.e. between branches (Palmer et al., 1991, Hansen 1997). Conversely, authors have suggested that branches can be considered as autonomous (Sprugel *et al.*, 1991). For instance, in shading experiments on walnut, sunlit branches have been observed to grow faster than shaded ones without any allocation of carbon to distant sinks (Lacointe *et al.*, 2004), thus emphasizing the sink strength limitation to long distance transport. This limitation of long distance carbon transport has potential impacts on developmental and growth processes. For instance, part-tree thinning of flower cluster has been shown to enhance branch vegetative growth and floral induction in the thinned tree sides (Palmer *et al.*, 1991; Grossman and Dejong, 1998). Similarly, shoot growth and to some extent starch accumulation in woody organs are impacted by fruit proximity (Berman and Dejong, 2003; Castillo-Llanque and Rapoport, 2011). Moreover, carbon transport and allocation change during a season. Indeed carbon labelling experiments have shown that carbon is allocated from reserves to support new shoot growth in spring (Kandiah, 1979a). At fall, carbon accumulate in leaves moves at long distances to roots to contribute to root growth and storage, before being reallocated to new growth in the next year (Hansen, 1967; Kandiah, 1979b).

In plant, carbon allocation between organs is commonly analyzed through the variations in organ biomass and non-structural carbohydrate (NSC) content. Among the different NSC forms in apple tree, starch, sorbitol and sucrose are the carbohydrates directly derived from the photosynthetic activity, with sorbitol and sucrose being the mobile forms for carbohydrate transport (Escobar-Gutiérrez 1996, Teo et al., 2006). Sorbitol and sucrose are transferred through the phloem to the sinks where they are converted into glucose and fructose (Teo *et al.*, 2006). Starch is commonly stored in reserve organs during the vegetative season. This NSC form is accumulated in reserve organs, where it can be mobilized for regrowth in spring or to buffer source-sink imbalances during the growing season (Sala *et al.*, 2012). Moreover, starch concentration, is directly associated to the ratio between source activity and sink demand (Naschitz *et al.*, 2010; Sala *et al.*, 2012).

In this study, we assumed that distances among organs and availability of resources are involved in both organ development (here floral induction) and growth (here considered as mean fruit weight). Our aim was to investigate the relative contribution of tree carbon balance, source-sink distances and GA availability in SAM on the FI and fruit growth. For this, we manipulated within-tree source-sink relationships during two years on ‘Golden delicious’ apple cultivar. We set trees in either high (ON trees) or low (OFF trees) crop loads, reducing by half the number of leaves (sources for both carbohydrates and florigen) or fruit (sources of gibberellins and sinks for carbohydrates). These manipulations were performed at different scales in the trees (shoots, branches and one side of the Y-shape trees) in order to clarify the effect of distances between sources and sinks. Moreover, we considered the within-tree variations in carbon acquisition (leaf photosynthetic activity) and accumulation (NSC concentration) and the gibberellins content in SAM to explore their respective involvement on FI. This study provides (i) new evidence of the likely co-involvement of gibberellins from fruits and signals originated from leaves other than carbohydrates in FI control and (ii) new elements on the debate on organ autonomy in trees with respect to carbohydrate transport in trees.

## Material and methods

### Plant material and growing conditions

The experiment was carried out from 2016 to 2018 on 10-year-old apple trees (cv. ‘Golden Delicious’). The orchard was located at the SudExpé experimental station in Marsillargues, in the south of France (43°66’N 4°18’E). Trees were initially planted in 1998 with ‘Tentation’ cultivar grafted on ‘Pajam2’ rootstock and then top grafted with ‘Golden delicious’ in 2005. The orchard was composed of four rows of 75 trees, each tree composed either of one vertical axis or of two main axes (Y-shape trees) arising five centimeters above the grafting point. Pruning and thinning were applied according to commercial practices before the beginning of the experiment in 2016. During the experimental period, all the trees were irrigated and fertilized to avoid any water or mineral deficiency.

### Varying source-sink relationships by leaf and fruit removal treatments

In spring 2016, around 65 trees in the orchard were set in OFF conditions by removing all the flowers after full bloom. No thinning was performed on the same number of trees, in which complete fruit removal was performed in 2015 to get high crop load in 2016. These trees were then considered as ON trees. In the following year (2017), the cropping status was reversed. In this years, all the flowers of OFF trees were removed just after full bloom to ensure a crop load equal to zero. To determine the period of FI in our conditions, a specific experiment was carried out. Assuming that the period of irreversible inhibition of FI was between 30 and 70 days after full bloom (DAFB) (Foster et al., 2003, Haberman et al., 2016), young fruit were completely removed on a selection of ON trees, at successive dates during this period. Fruit removal was performed at six dates (one tree per date): 30, 36, 42, 50, 56 and 70 DAFB in 2016 and at four dates in 2017 (37, 43, 50, and 58 DAFB).

On the two tree subsets in either ON or OFF conditions, 11 different treatments were set up (three trees per treatment) in springs 2016 and 2017. In order to modify fruit and leaves number, half of the leaves or half of the fruit were removed on the trees. Moreover, leaves and fruit were removed in different parts of the trees in order to modify the distances between the remaining leaves and fruit. Leaves and fruit were removed on either half of the shoots, half of the branches or one side of the Y-shape trees (Figure 1). New leaves that appeared after the first defoliation in spring were frequently removed throughout the growing season on the trees subjected to leaf removal. Trees not subjected to leaf or fruit removal either on ON and OFF crop load, were considered as controls. Another set of trees (called additional trees), in either ON or OFF conditions and not subjected to leaf or fruit removal was used to build a reference relationship between tree crop load, proportion of FI and mean fruit weight. 69, 103 and 65 trees of the field were considered in 2015, 2016 and 2017 to build this relationship.

**Figure 1.**
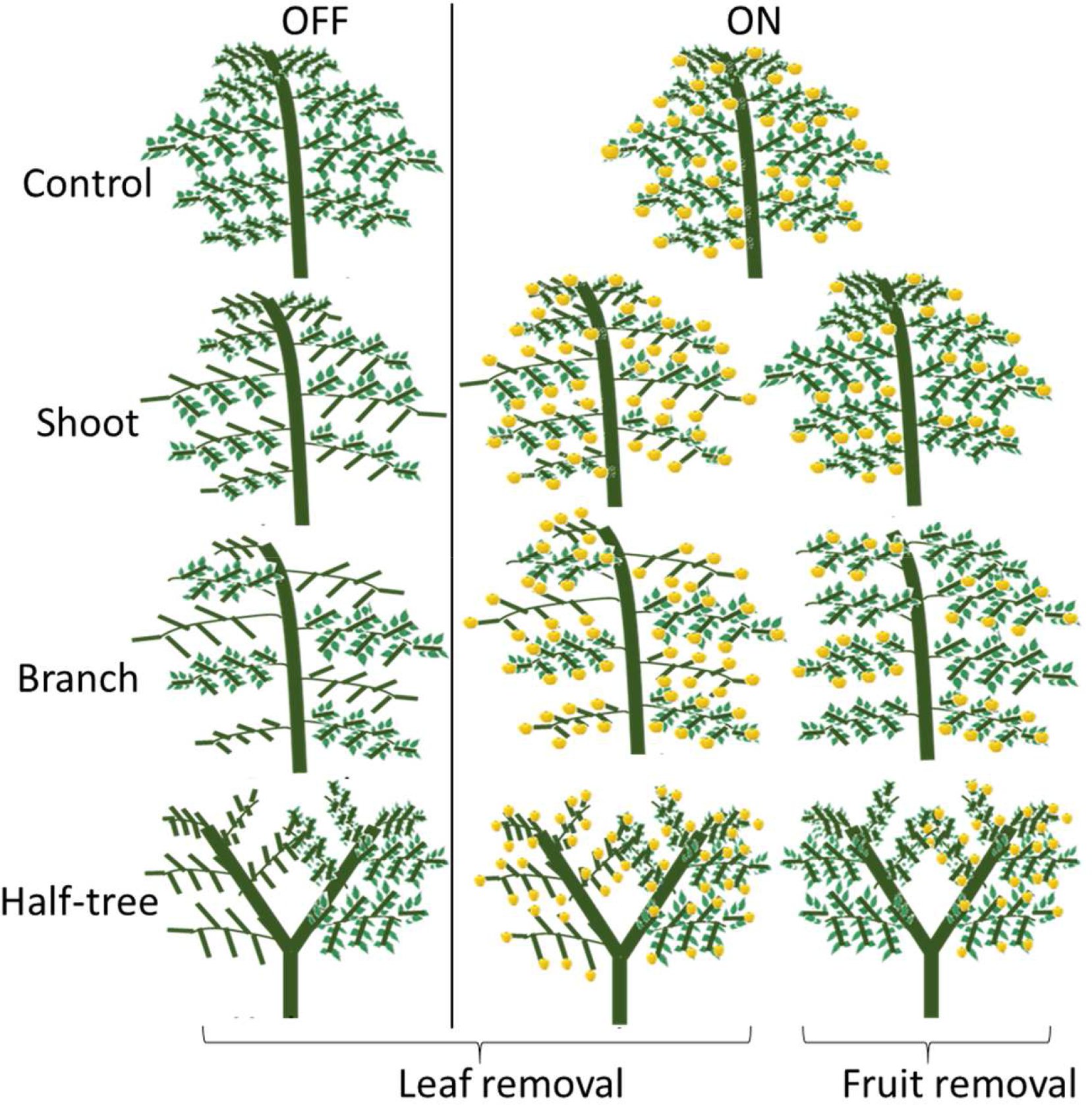
Schematic representation of the leaf and fruit removal treatments.

Crop load was estimated in each year by dividing the harvested fruit number per tree by its trunk cross sectional area (e.g. Francesconi et al., 1996). Trunk cross sectional area was computed assuming a cylinder shape by measuring in spring the trunk circumference at 10 cm above the grafting zone. For Y-shape trees, crop load was computed for both sides of the tree separately, considering them as mono-axial trees. The tree crop load of Y-Shape trees was then determined, considering this treatment as a combination of two mono-axial trees, by the mean crop load of the two sides of the trees.

### Development and growth variables: floral SAM proportion and mean fruit weight at harvest

The treatment effect on FI proportion in SAM was estimated at full bloom in the spring following treatment in 2017 and 2018 on all the trees including the additional trees and those subjected to sequential thinning in spring. FI proportion was estimated as the ratio of the total number of reproductive buds to the total number of growing buds. This proportion was estimated on six randomly distributed first-order branches per tree in each treatment, considering the leaf or fruit removal conditions (foliated/defoliated, fruiting/non-fruiting, 3 branches per condition). Unfortunately, no data were recorded for the trees subjected to fruit removal at the shoot scale in 2016.

At harvest, in early September of each year, fruit were collected on each treatment. Fruit were sorted by different parts of each tree considering whether they were subjected or not to leaf or fruit removal. All the fruit were collected on each tree except for the treatment performed at the shoot scale for which fruit were collected on two branches per tree, only. Then, each set of fruit was weighted and the mean fruit weight was estimated as the ratio of the total fruit weight to the number of fruit.

### Responses of leaf photosynthesis and starch content

Leaf photosynthesis and NSC contents were measured on August 2017 (from 119 to 145 DAFB) on fully expanded leaves belonging to short or medium shoots (shorter than 20cm, Costes et al., 2003) and fully exposed to sunlight. Measurements were performed on ON and OFF trees and on foliated parts of the trees with leaf removal treatments and on both fruiting and non-fruiting parts of the trees with fruit removal treatments. Three measurements were performed for each tree and condition (fruit or leaf presence/absence). Measurements were done between 8 and 12 am, with a infra-red gas analyzer (LI-6400, LICOR, Lincoln, Nebraska, USA) under controlled conditions within the growth chamber known to be non-limiting for photosynthesis (Massonnet *et al.*, 2007) (photosynthetic photon flux density = 1800 μmol^−2^s^−1^, relative humidity = RH = 70%, CO_2_ = 400ppm, T = 25°C).

After each photosynthesis measurement, the leaf, the entire annual shoot (called stem) on which the leaf was located, the SAM of this shoot and a 5 cm section of the one-year-old wood supporting it were sampled for measuring their NSC content. Three replicates of all these organs were sampled on each tree and for each condition (fruit or leaf presence/absence). Samples were placed immediately in liquid nitrogen and stored at −20°C for about one week. Then, they were freeze-dried and grinded to fine powders using a ball grinder. Starch concentrations were then determined for all the organs. In SAM, glucose, fructose, sorbitol and sucrose concentrations were also evaluated. All these analyses were performed following the protocol described in Pallas et al., (2018).

### GA concentrations in SAM

GA content measurements, were performed on SAM collected on 31 May 2017 (58 DAFB), i.e. at the expected date of FI in short shoots (Foster *et al.*, 2003). SAM were sampled on short to medium shoots that had recently stopped growing and did not formed protecting scars yet. SAM were collected on ON and OFF control trees, on trees subjected to fruit or leaf removal on half of the branches and on trees subjected to fruit removal on one side of Y-shape trees. Nine SAM were collected on each part of the trees (foliated/defoliated, fruiting/non-fruiting) and were gathered together for each tree. All samples were conserved at −80°C before being freeze dried and sent for GA quantification at the Plant Hormone Quantification Service in the Institute for Plant Molecular and Cell Biology (IBMCP), Valencia, Spain.

Fourteen GA forms produced in the two GAs biosynthesis pathways regulated by the activities of GA20-oxidases (GA20ox), GA3-oxidases (GA3ox) and GA2-oxidases (GA2ox) were investigated (Supplementary material figure S1). They include bio-active forms (GAs 4 and 1), degradation forms (GAs 51, 34, 29 and 8) and intermediate forms (GAs 12, 15, 24, 53, 44 and 19)

### Statistical analyses

All statistical analyses were performed with R software (R Development Core team, 2013). We investigated the effects of the combination of (i) the tree crop load status (ON or OFF trees), (ii) the tree treatment (control, leaf removal, and fruit removal), (iii) the scale (tree, shoot, branch, and one side of the Y-shape tree) at which treatments were performed and (iv) the condition within the tree (foliated, defoliated, fruiting and non-fruiting). The effect of all these combinations was tested on photosynthesis, NSC concentrations, GA concentrations and FI proportion, with a one-way ANOVA followed by a Tukey HSD test for pairwise comparison. Linear models were used for continuous variables and a general linear model of the binomial family was used for FI proportion. For GA and due to the low number of replicates (one per tree and condition), the effect of fruit presence/absence was also tested using Kruskal-Wallis test gathering samples from control ON and fruiting parts of trees (originated from branch and Y-Shape treatments) and from control OFF and non-fruiting parts of trees.

The dataset of additional trees with a large range of crop loads, obtained in 2015, 2016 and 2017 was used to fit a relationship between the tree crop load and the FI proportion (sigmoidal adjustment) or the mean fruit weight (exponential adjustment). The residuals between observed values for a given treatment and the general trend of FI proportion or fruit weight over different crop loads were used to test the treatment effects under comparable crop load conditions. As for raw variables, treatment effects on residuals were assessed by a one-way ANOVA followed by a Tukey HSD test.

## Results

### Tree crop load differed between treatments

Crop load was lower in 2016 and 2017, independently of treatments and displayed values equal to 12.1 and 20.7 fruit.cm^−2^ for control ON trees (Table 1). Crop load varied significantly among treatments when the treatments were compared altogether (P=0.0212 in 2016 and P=0.0012 in 2017). Considering values averaged over 2016 and 2017 and compared to control ON treatments, tree crop load was reduced by 25.6 %, 54.3% and 32.4% by fruit removal treatments at the shoot and branch scales and on one side of Y-shape tree, respectively. A lower crop load compared to the control ON trees was observed for all the leaf removal treatments except shoot and branch treatments in 2016 but this difference remained non-significant. Moreover, in each year, crop loads also varied among the three trees of each treatment, with large standard deviations observed in some cases.

**Table 1:**
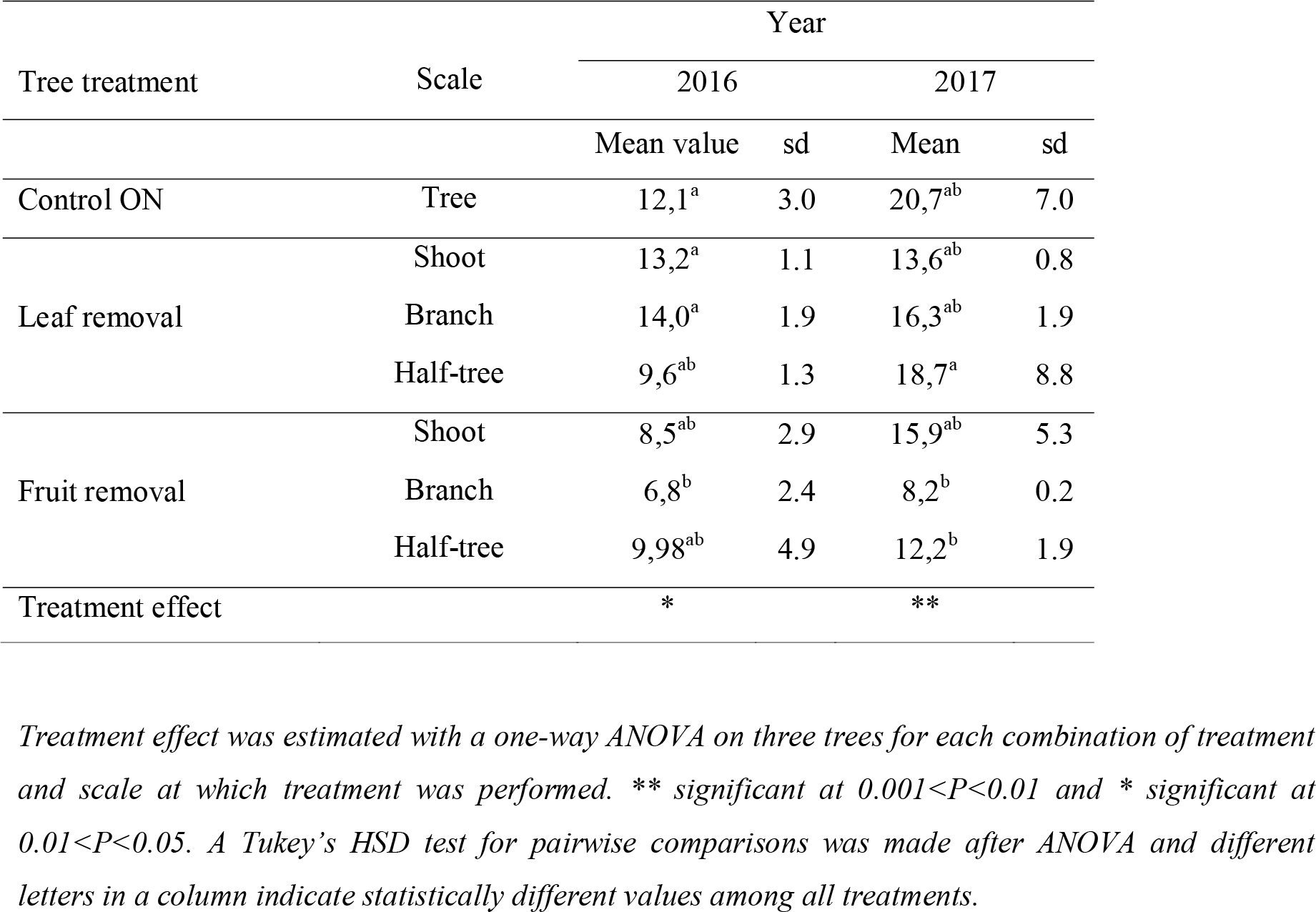
Mean and standard deviation (sd) values of crop load estimated as the fruit number per trunk cross sectional area (fruit.cm^−2^) for different treatments and scales at which treatments were performed for the control ON trees of ‘Golden delicious’ apple cultivar in 2016 and 2017.

### Floral induction in SAM occurs after treatment onset

The complete fruit removal performed sequentially in springs 2016 and 2017 on a subset of ON trees allowed evaluating the date after which the inhibition of FI by fruit presence was no longer reversible. The quantification of FI proportion in the following spring revealed that FI was no longer possible at 70 DAFB (Table 2). At that date, FI proportion reached values similar to those of control ON trees. Conversely, when fruit removal was performed before 50 DAFB, FI proportion was close to 100% as observed for OFF control trees, in both years (Table 2). Assuming that dates of FI were similar for all the buds in trees, this suggests that FI likely occurred during a short period between 50 and 70 DAFB. This shows that our experimental design was relevant since treatments were performed before 50 DAFB at a date when the SAM fate was not yet determined.

**Table 2:** Proportion of shoot apical meristems (SAM) induced to flower in ‘Golden delicious’ apple trees subjected to complete fruit removal performed sequentially from 30 to 70 days after full bloom (DAFB), in 2016 and 2017 and for control OFF and ON trees. FI proportions were evaluated based on the floral bud proportion, as the ratio of the total number of reproductive buds to the total number of growing buds, on six branches per tree, in the next spring, after the year of treatment.

| Treatment | Floral induction 2017 |  | Floral induction 2018 |  |
| --- | --- | --- | --- | --- |
|  | Date of fruit removal 2016 | FI proportion | Date of fruit removal 2017 | FI proportion |
|  | (DAFB) |  | (DAFB) |  |
| Control OFF | - | 97% | - | 100% |
| Fruit removal | 30 | 95% | - | - |
|  | 36 | 79% | 37 | 98% |
|  | 42 | 70% | 43 | 100% |
|  | 50 | 76% | 50 | 90% |
|  | 56 | 40% | 58 | 56% |
|  | 70 | 18% | - | - |
| Control ON | - | 19% | - | 29% |

### FI proportion is affected by leaf and fruit presence and by their distance to the SAM

During the experiment and on all the additional control trees, FI proportion was strongly associated with the tree crop load in the previous year. The relationship between FI and tree crop load was fitted with a logistic decreasing function (Figure 2 and supplementary material Figure S2).

**Figure 2.**
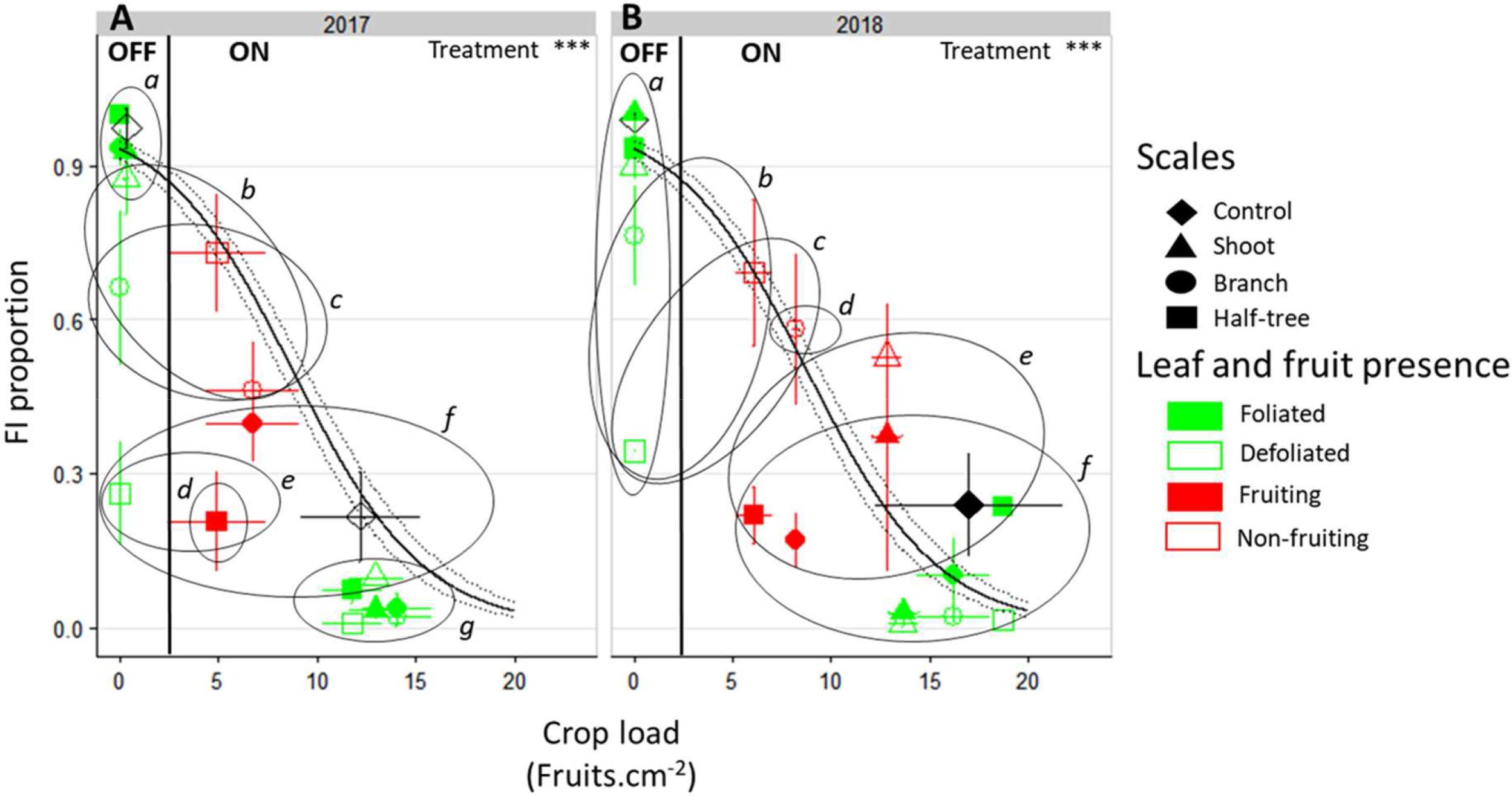
Relationship between FI proportion and tree crop load (number of fruit per trunk cross sectional area) in ON and OFF ‘Golden Delicious’ apple trees for the different treatments in 2017 (A) and 2018 (B). Each point represents the value for one combination of tree treatments, tree scale at which treatments were performed and conditions within the trees (leaf or fruit presence) and bars represent the standard deviation among measurements (3 measurements for each treatment combination). The continuous line represents the logistic function fitted on the additional tree dataset (y = exp (−0.3008 × x + 2.6341) / (1 + exp (−0.3008 × x + 2.6341))) (see supplementary material S2). The dotted grey lines represent the deviation interval of the fitted values. The dataset was fitted with a glm model (binomial family) and leaf and fruit presence effect was assessed with one-way-ANOVA. *** significant at P<0.001. A Tukey’s HSD test for pairwise comparisons was made after the analysis and different letters indicate statistically different values among all conditions.

Leaf removal did not impact FI on ON trees, with values close to zero on both foliated and defoliated parts, whatever the scale at which leaf removal was performed (Figure 2, right side). In the foliated parts of the defoliated OFF trees FI proportion was similar to the control OFF trees (between 0.9 and 1, Figure 2, left side). In contrast, a strong and significant decrease in FI proportion was observed in the defoliated compared to the foliated parts of trees after leaf removal on half of the branches (−29% and −19% for 2017 and 2018 respectively) or half of the tree (−74% and −63% for 2017 and 2018 respectively). Stronger decrease in the defoliated side of Y-shape trees scale than on defoliated branches was observed, suggesting an impact of the distances to the remaining leaves on FI. This distance effect was also found by the absence of any significant decrease in FI on the defoliated shoots (−6% and −10% for 2017 and 2018 respectively) of trees subjected to leaf removal at the shoot scale (Figure 2). Local fruit removal had also a strong effect on FI proportion within the tree (Figure 2 A and B, right side). FI proportions were lower in the fruiting than in the non-fruiting parts. Consistently with the distance effect observed after leaf removal treatments, FI proportion increased in non-fruiting parts compared to the fruiting ones when the distances to the remaining fruit increased. In 2018, the increase in FI proportion between fruiting and non fruiting parts was equal to 50.1% for Y-shape trees while it only reached 29% when fruit removal was performed at the shoot scale.

Residual values of FI proportion estimated from the logistic function (supplementary material Figure S2) were used to analyze the treatment effects with respect to the values predicted for trees under similar crop load (Table 3). This analysis allowed us to inspect the effect of fruit removal treatments that directly modified the tree crop load. After fruit removal treatments, the residuals of FI proportion in spring 2017 and 2018 were significantly lower in the fruiting parts of the trees than the values predicted under similar crop load (Table 3) for fruit removal treatments at the branch scale and one side of the Y-shape tree. In contrast, FI proportion in the fruiting and non-fruiting parts was significantly higher than the predicted values for trees with similar crop load when fruit removal was performed at the shoot scale. Both together these results suggested that FI could be directly driven by the crop load at the tree scale but only when fruit load distribution within the tree is homogeneous (after fruit removal at the shoot scale).

**Table 3:**
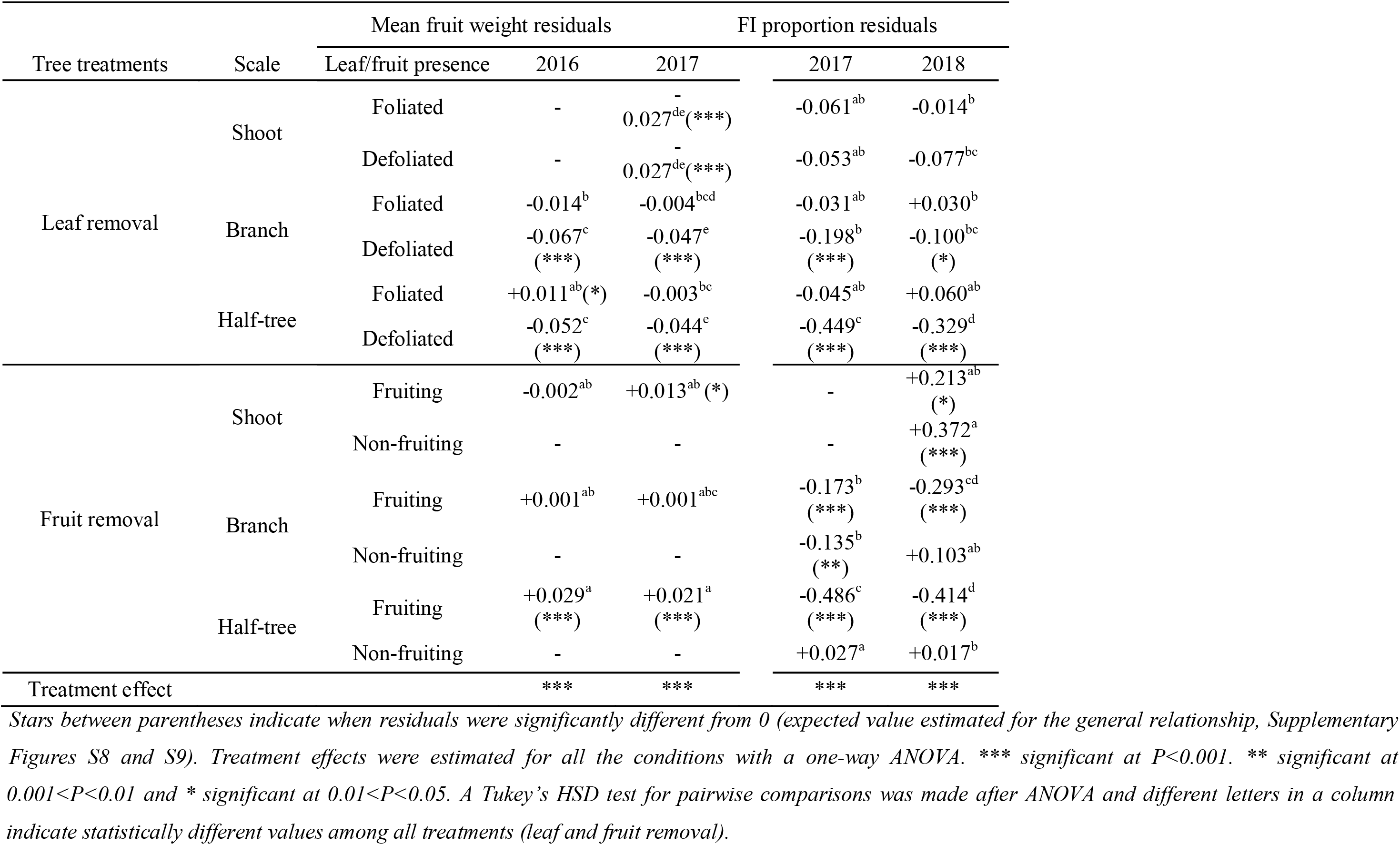
Mean values of residuals extracted from non-linear adjustments between the mean fruit weight (kg) or FI proportion and tree crop loads on ‘Golden delicious’ apple trees. Data are presented for different treatments and scales at which treatments were performed and depending on leaf or fruit removal conditions.

### Mean fruit weight is affected by distances between leaves and fruit

The general trend of mean fruit weight over crop loads, for the additional control trees over three years, displayed a negative relationship (Figure 3 and supplementary material Figure S3) with highest mean fruit weights equal to around 0.25kg and lowest ones to 0.08kg.

**Figure 3.**
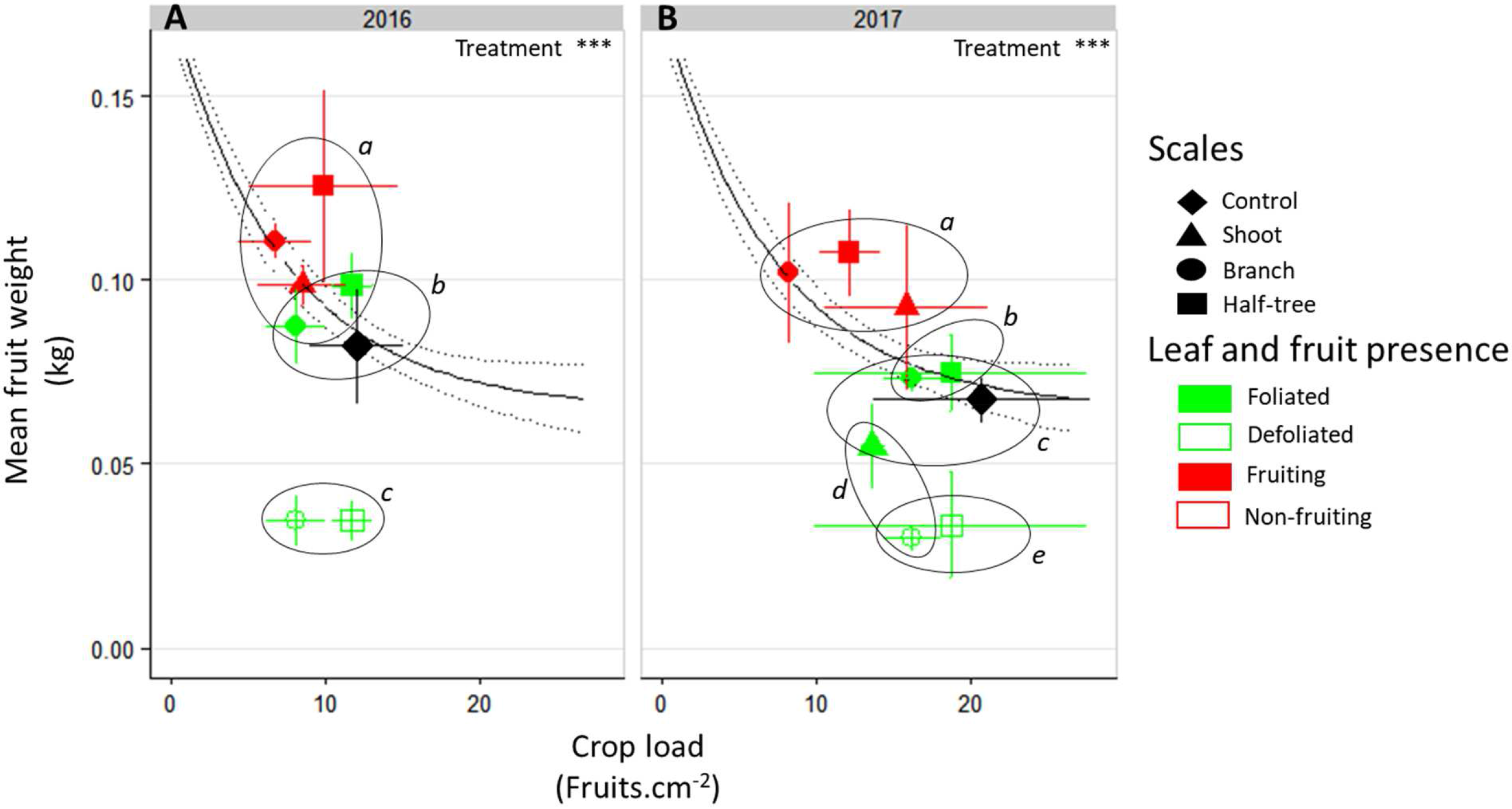
Relationship between mean fruit weight and crop load in ON ‘Golden Delicious’ apple trees for the different treatment combinations in 2016 (A) and 2017 (B). Each point represents the value for one combination of tree treatments, tree scale at which treatments were performed and condition within the trees (leaf or fruit presence) and bars represent the standard deviation among measurements (3 measurements for each treatment combination). The continuous black line represents the exponential function fitted on the additional trees dataset (Y= 0.5 × exp (−0.2 × x) + 0.06) (supplementary material Figure S3). The dotted grey lines represent the deviation interval of the fitted values. Leaf and fruit presence effect was estimated with a one-way-ANOVA. *** significant at P<0.001. A Tukey’s HSD test for pairwise comparisons was made after the analysis and different letters indicate statistically different values among all combinations.

As for FI, mean fruit weight depended on the distances to remaining leaves in trees subjected to leaf removal (Figure 3). Indeed, mean fruit weight decreased of 53% and 59% in the defoliated parts of trees subjected to leaf removal at the branch scale or on half of Y-Shape trees, respectively compared to the foliated parts. Conversely, this decrease was no longer significant (equal to 2%) when defoliation was performed at the shoot scale. In both foliated and defoliated parts of these trees (defoliation at the shoot scale), mean fruit weight was not significantly different to what observed for control ON trees. Fruit removal increased mean fruit weight compared to control ON trees. Nevertheless, this increase was higher when fruit removal was performed at the branch scale or on one side of Y-Shape trees than after fruit removal at the shoot scale.

The analysis of residuals to relationship between crop load and mean fruit weight (Table 3) showed that the mean fruit weight in the fruiting shoots (2016) and branches (2016 and 2017) after fruit removal treatments was similar to that of control trees with similar crop load and with a homogeneous distribution of fruit within the tree. A slight increase in residual values (around +0.02 kg) was observed when fruit removal was performed on one side of the Y shape trees in both years or on half of the shoots in 2017. These results suggest that fruit weight was mainly determined by the tree crop load whatever the distance to remaining fruits after fruit removal.

### Relationships between FI and mean fruit weight variability and carbon availability

Photosynthesis rate was higher for ON trees compared to OFF ones (mean values 7.6 and 14.3 μmol.m^−2^.s^−1^ for OFF and ON control trees, respectively; Figure 4). Moreover, no significant difference in photosynthetic rates (mean value 15.76 μmol.m^−2^.s^−1^ for all fruit removal treatment) was observed between the different fruiting and non-fruiting parts of trees subjected to fruit removal. This suggests that fruit presence stimulated photosynthesis whatever the distances to the fruit.

**Figure 4.**
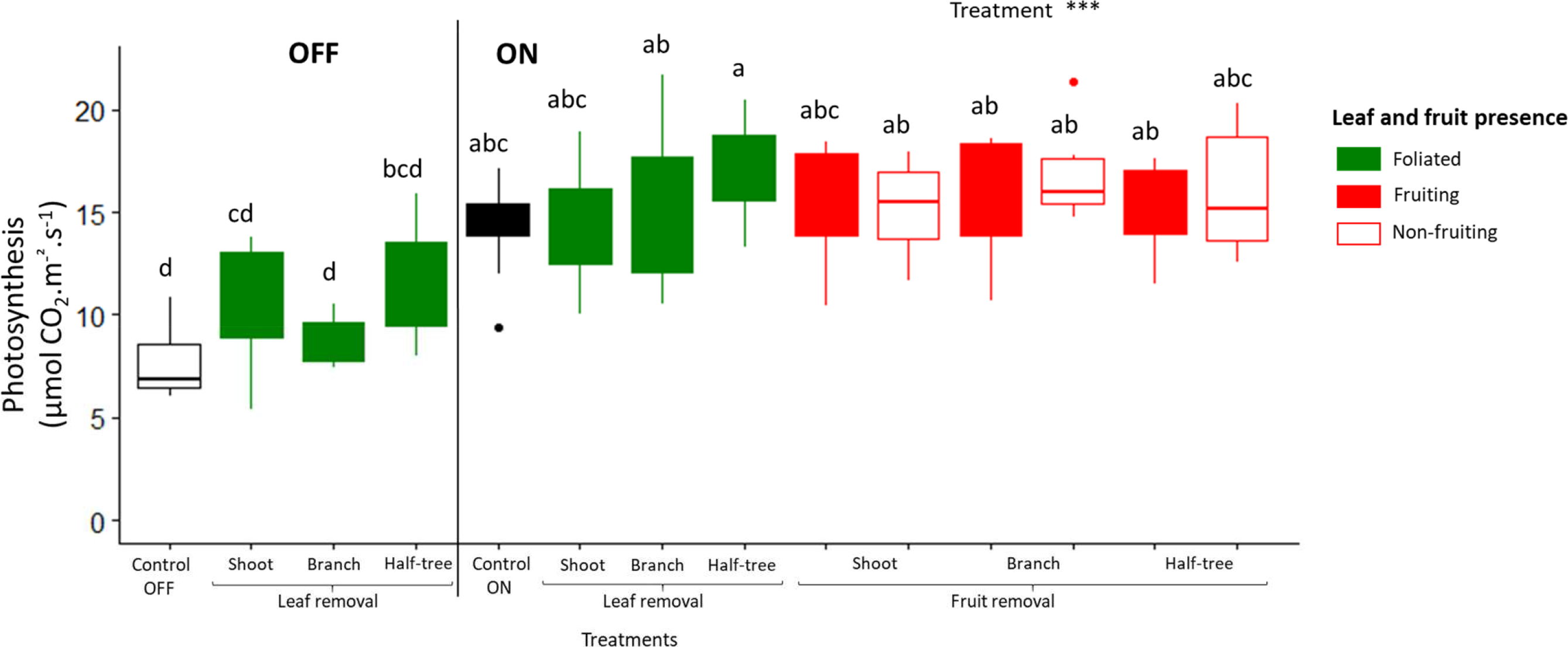
Boxplot representation of leaf photosynthetic activity in August 2017 for ‘Golden Delicious’ apple trees for the different treatments (control, leaf removal, and fruit removal), tree scales (tree, shoot, branch, and one side of Y-shape trees) at which treatments were performed and conditions within the tree (foliated, defoliated, fruiting and non-fruiting) for ON and OFF trees. Nine replicates were used for each treatment combinations (3 samples × 3 trees). Treatment effect was estimated with a one-way-ANOVA considering all the combinations together. *** significant at P<0.001. A Tukey’s HSD test for pairwise comparisons was made after the analysis and different letters indicate statistically different values.

Starch concentration varied among organs, with low values in SAM and leaves (Figure 5 A, B) and higher values in stems and wood (Figure 5 C, D). Tree crop load negatively affected starch concentration in leaves, stem and wood but had no effect on starch concentration in SAM. (Figure 5 B) with similar values between ON and OFF trees. In OFF trees, no impact of defoliation treatments was observed on starch concentrations in SAM, stems and wood (Figure 5, left sides), although a decrease in FI proportion was observed in defoliated parts of OFF trees. In contrast, in ON trees, significantly higher starch concentration were found in the SAM, stems and wood when comparing the leafy parts to defoliated ones (Figure 5 B, C, D). For these trees, this effect of leaf removal was similar regardless of the distances to the remaining leaves since no difference was observed between defoliation at the shoot, branch or half tree scale. The effect of fruit removal on starch concentration was observed in wood only, through a greater concentration in the non-fruiting than in the fruiting parts of defoliated shoot, branch and one side of the Y-shape tree treatments (Figure 5D). No clear impact of the distances to the remaining fruit was observed on starch concentrations in wood, as the decrease in starch content were similar whatever the scale at which fruit removal was performed.

**Figure 5.**
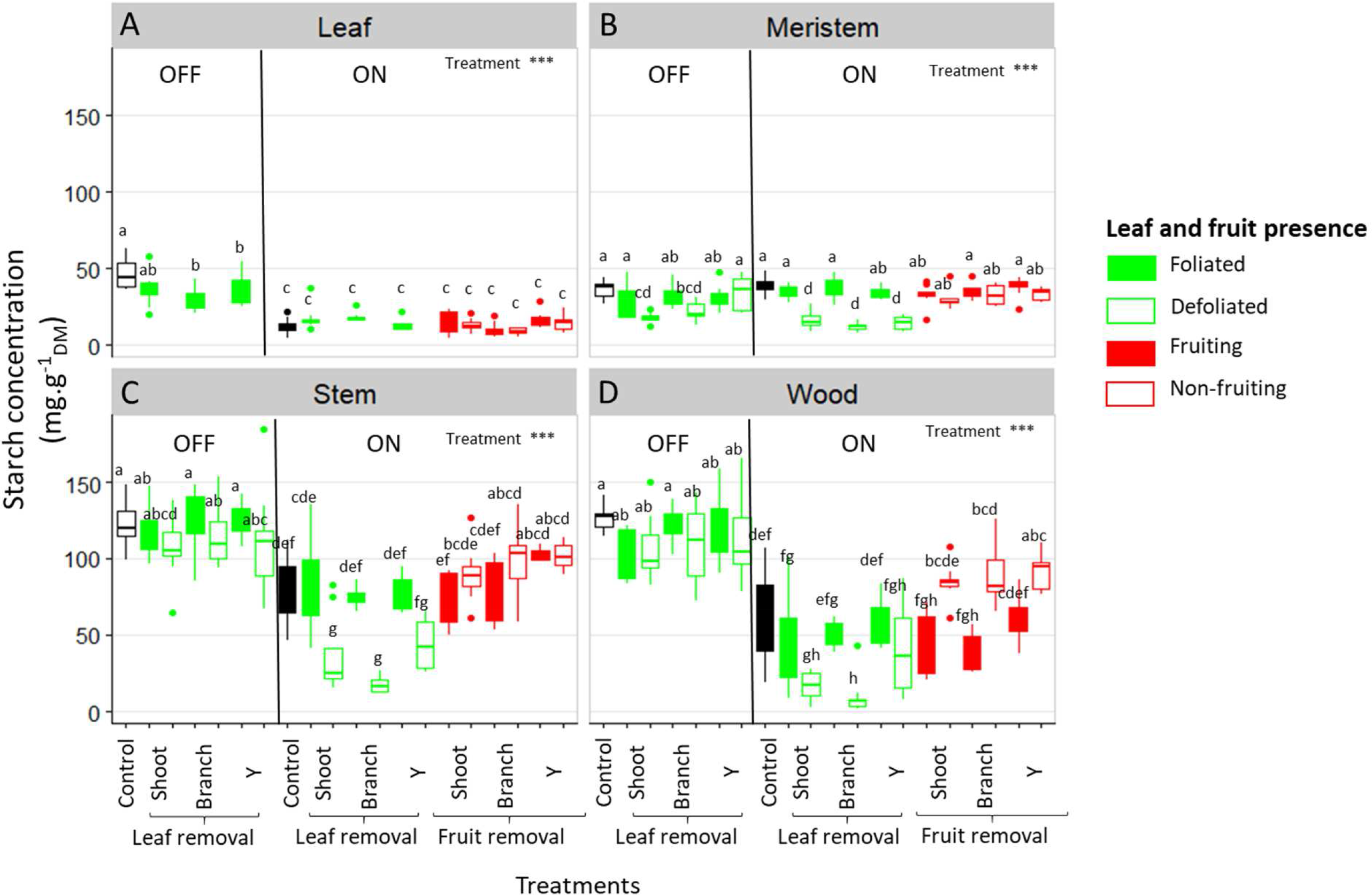
Boxplot representation of starch concentration in the leaves (A), shoot apical meristems (B), stems (C) and one-year-old wood (D) of ON and OFF ‘Golden Delicious’ apple trees for the different treatments, tree scales at which treatments were performed and conditions within the trees (leaf or fruit presence). Nine replicates were used for each treatment combinations (3 samples × 3 trees). Treatment effect was estimated with a one-way-ANOVA considering all the combinations together. *** significant at P<0.001. A Tukey’s HSD test for pairwise comparisons was made after the analysis and different letters indicate statistically different values among all treatments.

Regarding, the soluble sugars content (sorbitol, sucrose, fructose and glucose) in SAM (supplementary material Figure 4), sorbitol displayed higher concentrations than the three other sugars. Moreover, treatment effects (fruit and leaf removal) on soluble sugars content in SAM were observed on sorbitol concentration, only after leaf removal treatments on both OFF and ON trees.

### GA9 (precursor of active form) and GA1 (active form) decrease in fruit presence

In the early-13-hydroxylating pathway (supplementary material Figures S5 and S6), three forms were found in abundance (Table 4): GA44 (inactive form), GA1 (active form), and GA8 (degradation form). In the non-hydroxylating GA pathway, maximal GA concentrations were found for GA9 (Table 4) which is the last inactive form before GA4 synthesis. Variations in SAM GA contents were observed among all the sampled trees (supplementary material Figure S6). Nevertheless, differences were significant between SAM from fruiting and non-fruiting parts of trees. GA9 concentration was significantly higher in the SAM collected on the non-fruiting trees or parts of trees (gathering control OFF trees and in non-fruiting branches and sides of Y-shape trees) than in those collected on fruiting trees or parts of trees (gathering control ON trees, fruiting branches and sides of Y-shape trees) (Table 4). Even though not significant, higher concentration was observed in the non-fruiting side of the Y-shape tree than in the non-fruiting branches, suggesting a possible effect of distances to the remaining fruit. In addition, a slightly higher but non-significant GA1 concentration was observed in the SAM of non-fruiting parts of the trees than in the fruiting ones. Conversely, GA8 concentration was higher in control ON than in control OFF trees and in fruiting than in non-fruiting branches when fruit removal was performed on half of the branches. Nevertheless, no difference between fruiting and non-fruiting branches was observed for the Y-Shape trees. Finally, leaf removal did not influence any GA concentration.

**Table 4:**
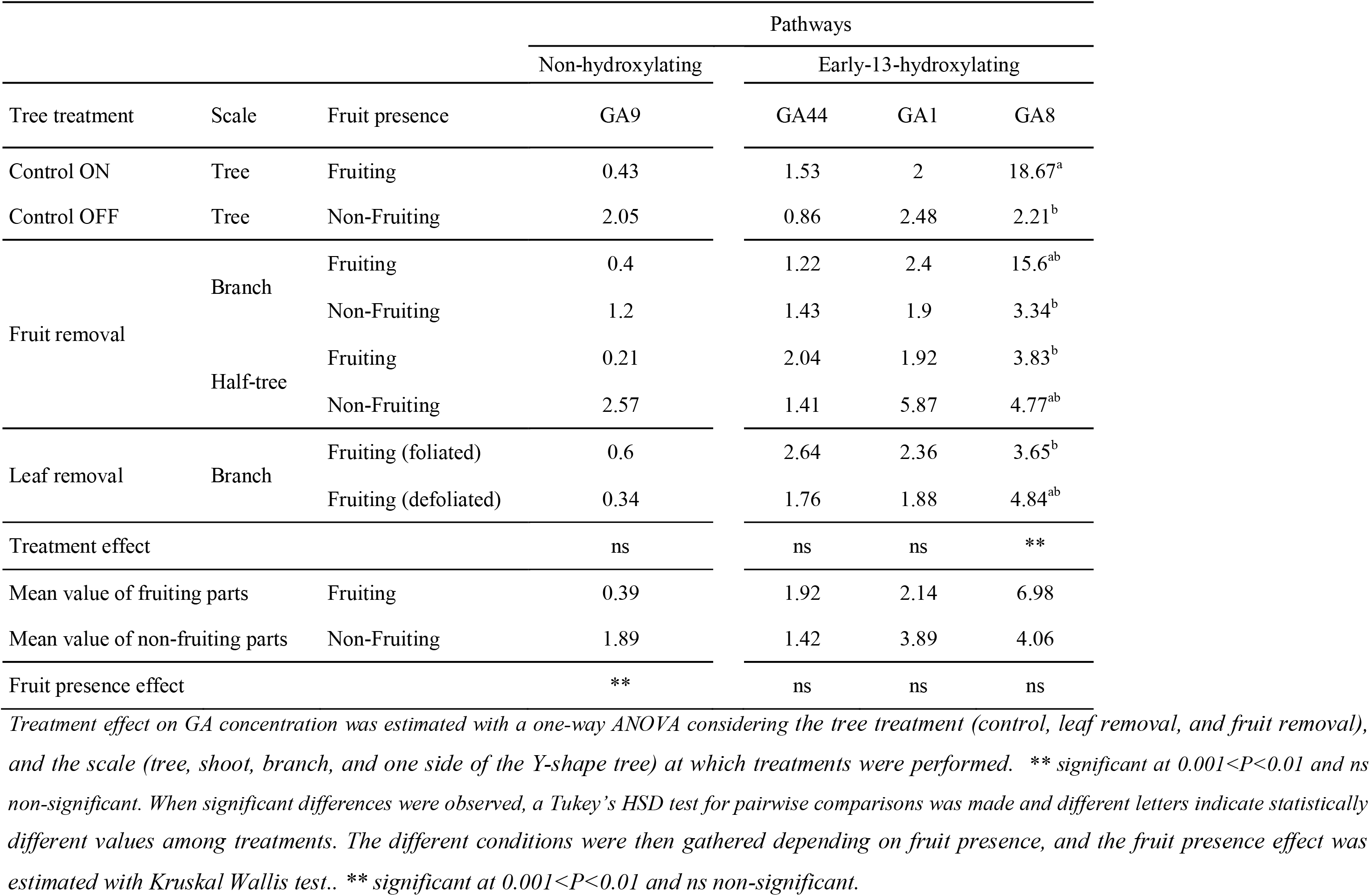
Concentations (ng.g^−1^) of GA9, 44, 1 and 8 in shoot apical meristems sampled at 58 days after full bloom on ‘Golden Delicious’ apple trees under different treatments (leaf or fruit removal), performed at different scales (branch, half-tree).

## Discussion

### Relative roles of carbohydrates and GA in flower induction

Our study investigated the impact of the tree carbon balance on floral induction by exploring the relation between NSC contents in all the organs and SAM status (floral induced or not) after organ manipulations (leaf or fruit removal). After defoliation treatments, the decrease in FI proportion in the defoliated branches and half-side of OFF trees was not associated with any decrease in starch content in all organs including SAM (Figures 5). In addition, NSC concentration, whatever the forms (soluble or starch), did not vary between fruiting and non-fruiting parts in all the organs including SAM (Figures 5 and S4), while a decrease in FI was observed in fruiting parts compared to non-fruiting parts. Together these results on trees subjected to leaf or fruit removal suggest that FI is not only related to the tree carbon balance and carbohydrate availability in SAM.

Other possible effects of leaf removal independently of the carbon production and primary metabolism (here analyzed through NSC and starch content) could be implicated. Indeed, leaves are likely to be sources of FT protein, considered as the florigen which is transported to SAM where it activates flowering (Corbesier and Coupland, 2006; Hanke *et al.*, 2007). Nevertheless, other carbohydrates than starch and NSC may have role in FI, especially signaling molecules such as trehalose-6-phosphate (T6P, Ponnu et al., 2011; Lastdrager et al., 2014). In Arabidopsis T6P has been shown to affect flowering, by inducing FT production and the expression of flowering-time (SPL) genes in SAM that in turn regulate flowering as a function of plant age (Wahl *et al.*, 2013).

Moreover, GA4 has been shown to target key flowering genes in SAM, in *Arabidopsis* (Eriksson et al., 2006). In apple tree, GA4 and GA1 may inhibit FI (Ramírez et al., 2001; Ramírez et al., 2004). In the present study, the inactive GA9 preceding GA4 in the non-hydroxylating pathway accumulated in OFF trees, and in non-fruiting branches and side of Y-shape trees (supplementary material Figure S5). Nevertheless, no difference in GA4 concentrations was found in SAM between fruiting and non-fruiting conditions. In addition, in the hydroxylating pathway, GA1 was slightly higher in non-fruiting tree parts and in OFF trees (supplementary material Figure S5) whereas GA8 slightly accumulated in fruit presence. Altogether these results are consistent with the down-regulation of *MdGA2ox* transcripts in OFF trees (or its up-regulation in ON trees), observed in Guitton et al., (2016). They suggest that the last steps of GA catabolism could be less active in absence of fruit (conversely more active in presence of fruit). Therefore, the putative role of GA on controlling FI is supported by our results and previous findings even though their inhibitory or activating effect remains to be clarified. Moreover, further researches would be needed to investigate the ability of GAs, likely produced by seeds (Dennis and Nitsch, 1966), to directly act on SAM FI in the apple tree.

### Response of floral induction and fruit growth to changing source-sink distances

In this study, leaf and fruit removal at different scales of plant organization allowed us to clarify the impact of distances between organs on FI and fruit growth. Similar FI proportion was observed between foliated and defoliated shoots and between fruiting and non-fruiting ones when leaf or fruit removal was performed at the shoot scale, thus implying transport at short distances of signals originated from leaves (activators) and fruits (inhibitors). This suggests that shoots can be considered as non-autonomous and prone to exchanges of inhibiting/activating signals.

In contrast, leaf presence in the foliated parts of trees subjected to defoliation at the branch scale or on one side of Y-Shape trees did not promoted FI in the defoliated parts of these trees. In that case, distances were too long, possibly for florigen transport, consistently with previous studies having underlined the lack of evidence of FT long distance transport in woody plants (Putterill and Varkonyi-Gasic, 2016). Similarly, FI was only slightly affected by fruit presence at long distance since FI in the non-fruiting branches or sides of Y-Shape trees was slightly lower or even similar to that observed on OFF trees. This suggests that the inhibiting signal produced by fruits may not be transported at long distances in the tree structure or in low quantity, only. GA transport at relatively long distances has been demonstrated in small annual plants, with GA20 being the mobile form in *Pisum sativum* (Binenbaum *et al.*, 2018) and GA12 the form transported through the xylem in *Arabidopsis thaliana* (Regnault *et al.*, 2015). Interestingly, the different GA forms issuing from the hydroxylated and non-hydroxylated pathways may involve different transporters (Binenbaum *et al.*, 2018). However, studies on the transport of these forms and the distances at which they could be transported in more complex plants such as in fruit trees are still needed.

Fruit weight was also strongly affected by the distances to the remaining leaves after leaf removal. As for FI, similar fruit weights were observed between neighboring leafy and non-leafy shoots, when defoliations were performed at shoot scale. This is consistent with previous studies on peach where non-fruiting shoots contributed to fruit growth in nearby fruiting shoots (Walcroft *et al.*, 2004). Conversely, a strong decrease in fruit weight and starch concentrations in all organs were observed on defoliated branches or defoliated parts of Y-Shape trees. This is in accordance with previous studies of carbon labeling on young walnut and peach trees, that have shown limitation of carbon transport at long distance leading to almost complete autonomy of branches even when exposed to source limitation through shading, leaf removal or girdling (Lacointe *et al.*, 2004; Volpe *et al.*, 2008). These results are also consistent with leaf removal effect on fruit growth and reserve accumulation in young fuyu trees and mature *Carpinus*, *Fagus* and *Tilia* forest trees (Choi *et al.*, 2003; Hoch, 2005).

Conversely, long distance transport of carbohydrates was suggested by the results after fruit removal. Indeed, mean fruit weight in the fruiting parts of trees subjected to fruit removal was similar or even higher to that observed in control trees with a homogeneous crop load (Table 3). Moreover, starch concentration in stems and leaves of non-fruiting parts of ON trees (Figure 5D) was lower than that observed in OFF trees suggesting carbohydrate export to the fruiting parts of the trees even at long distances to sustain fruit growth. Nevertheless, it is noticeable that a part of the carbon excess produced in the non-fruiting parts was allocated to the reserve organs. This confirms the major role of reserves as an active sink in perennial tress (Silpi *et al.*, 2007). The low NSC content observed in leaves of the non-fruiting parts can be interpreted as resulting from carbon export and prevented photosynthesis inhibition by starch accumulation (Wunsche *et al.*, 2000), thus leading to a similar photosynthesis rate in fruiting and non-fruiting parts of the trees (Figure 4).

A discrepancy on the distance effects on fruit growth between trees subjected to leaf and fruit removal may appear from our results. However, this apparent discrepancy results from the nature of each treatment. First, leaf removal likely affected the transpiration flux (e.g. Pataki et al., 1998) and may have disturbed the long distance transport of carbohydrates (e.g. Hölttä et al., 2009, Nikinmaa et al., 2013). Second, large within tree source-sink imbalances existed in the trees subjected to fruit removal with the non-fruiting parts displaying low sink demand and large carbohydrate supply and the fruiting parts displaying high sink demand. This imbalance could be the driver of carbon fluxes even at long distance while in the trees subjected to leaf removal fruit sink demand remained high in all the tree parts thus limiting carbon fluxes from the remaining leaves to the distant fruit (Walcroft *et al.*, 2004).

## Conclusion

Our results shows that SAM floral induction is not directly associated to the tree carbon balance nor organ starch content and NSC availability in SAM but more probably to the combination of activating and inhibiting signals originated from leaves and fruit. Having performed leaf and fruit removal at different scales of tree organization provides new clues for understanding the distances at which these signal can act within the plant. At short distances (neighboring shoots), these signals are able to move from sources (leaves and fruit) to sinks (SAM) to act on FI while they cannot reach SAM at longer distances (branches and sides of Y-shape trees). Moreover, this study suggests that carbohydrates can move at longer distances from branch to branch in condition of high source-sink imbalances within the tree and in absence of any perturbation of the vascular fluxes. Finally, this study brings new considerations on carbohydrate and hormone transports within the fruit trees that can be then integrated in functional structural plant model (e.g Vos *et al.*, 2009) to simulate floral induction and fruit growth over years.

## Aknowledgments

The authors thank the AFEF team, SudExpé for technical support during experiments, UMR PIAF for having provided LICOR 6400 device, CIRAD biochemistry platform for NSC analyses and IBMCP for GA quantification. The authors also thanks Dr. B. Guitton for his implication in the gibberellins results analysis. This work was funded by the ANR-DFG Alternapp project and by the PhD grant accorded to F. Belhassine by INRA BAP department and ITK.

## Supplementary material

**Figure S1.** Representation of the gibberellins biosynthesis pathways.

**Figure S2.** Relationship between crop load and FI proportion for the additional control trees.

**Figure S3.** Relationship between crop load and mean fruit weight for the additional control trees.

**Figure S4.** Sorbitol, sucrose, fructose and glucose concentrations in shoot apical meristems.

**Figure S5.** Boxplot representation of concentrations of all gibberellin forms (ng.g^−1^) in shoot apical meristems.

**Figure S6.** Distribution of concentrations of the GA9, 44, 1 and 8 in the two biosynthesis pathways in shoot apical meristems.

